## Supplementary Tabel and Figures for "A Common Pathway for Detergent-Assisted Oligomerization of Aβ42"

### Stabilization of an Aβ42 Tetramer in Detergent Micelles

*Supplementary Information*

Table S1. Information for the systems studied by molecular dynamics simulations

| Peptide | Parallel or antiparallel | Dimer or tetramer | SDS or DPC | N or C-sheet | # of detergents | Simulation time /replicate[a] ($\mu$s) |
|---|---|---|---|---|---|---|
| Aβ42 | parallel | dimer | SDS | N | 80 | 1 |
| Aβ42 | parallel | tetramer | SDS | N | 74 | 1 |
| Aβ42 | parallel | dimer | SDS | C | 59 | 1 |
| Aβ42 | antiparallel | dimer | SDS | C | 63 | 1.2 |
| Aβ40 | parallel | dimer | SDS | C | 62 | 1 |
| Aβ42 | parallel | tetramer | SDS | C | 62 | 2 |
| Aβ40 | parallel | tetramer | SDS | C | 61 | 1.1 |
| Aβ42 | antiparallel | tetramer | SDS | C | 70 | 2.2 |
| Aβ42 | antiparallel | tetramer | SDS | C | 61 | 1 |
| Aβ42 | antiparallel | tetramer | DPC | C | 71 | 1.5 (2.0[b]) |
| Aβ42 | antiparallel | tetramer | DPC | C | 61 | 1 (1.8[b]) |

[a]Four replicate simulations were run for each system.

[b]One of the replicate simulations was extended to a longer time.

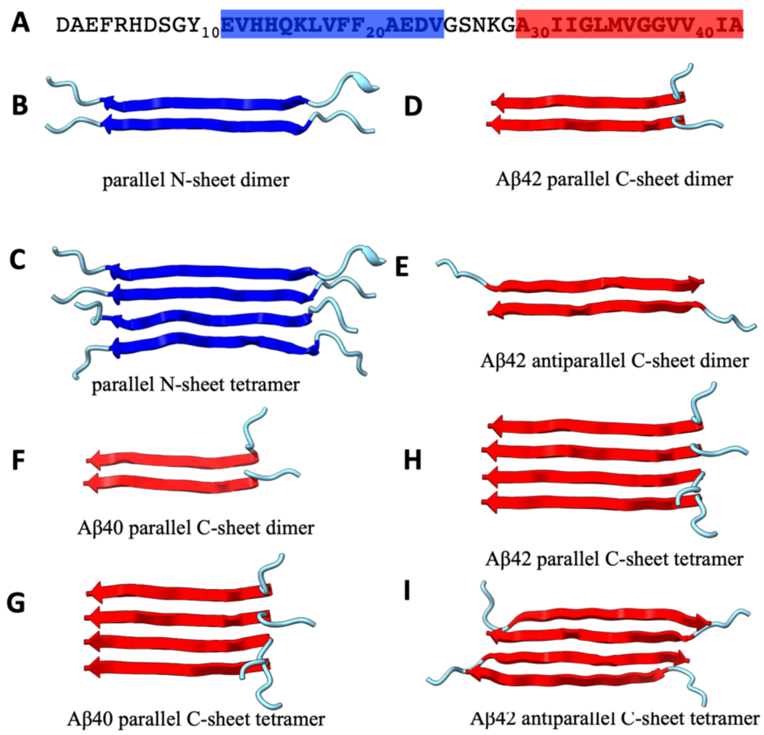


Figure S1. Aβ sequence and oligomers studied here. (A) Aβ42 sequence, with N and C-strand highlighted in blue and red respectively. (B-I) Dimeric or tetrameric parallel or antiparallel N or C-sheet.

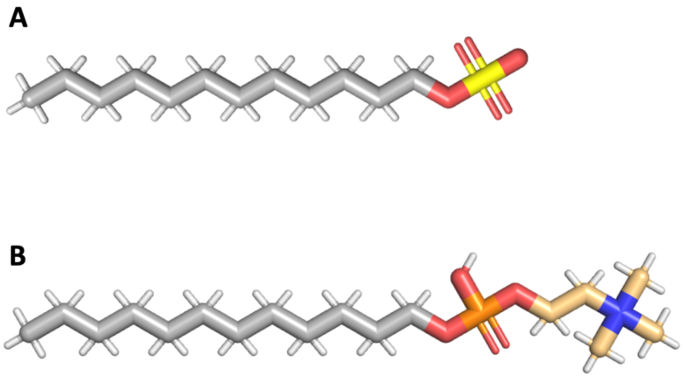


Figure S2. SDS and DPC molecules have the same hydrocarbon tail but different head groups. (A) SDS molecule. (B) DPC molecule. The colors of headgroup atoms are: S, yellow; P, orange, O, red, N, blue, C, light orange.

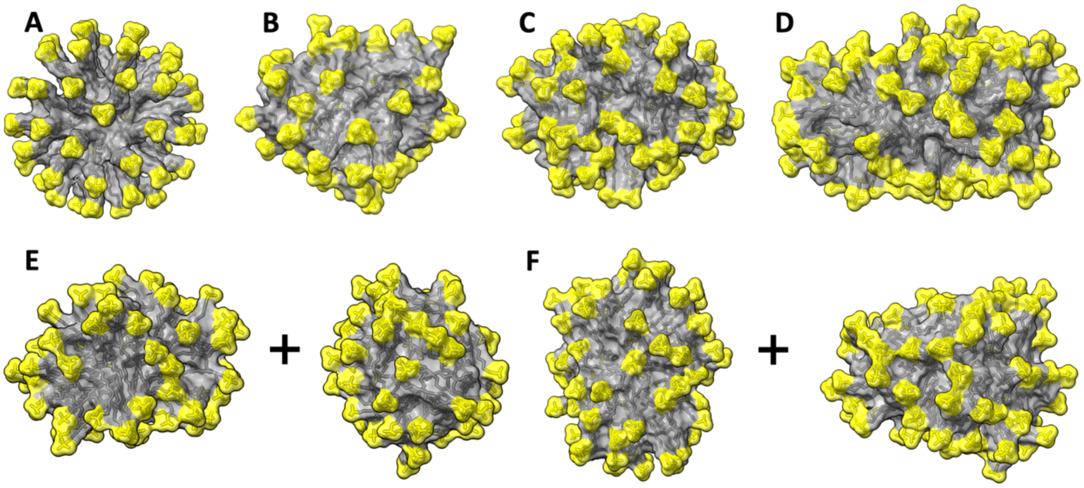


Figure S3. MD simulations of pure SDS micelles. (A) Initial structure of a micelle with 60 SDS molecules. (B-F) Final structures of micelles with 60, 80, 100, 120, and (F) 150 SDS molecules. In the last two cases, the initial micelle broke into two separate micelles. Each simulation was run for 500 ns. Micelles are rendered in surface representation, with headgroups in yellow and hydrocarbon tails in grey.

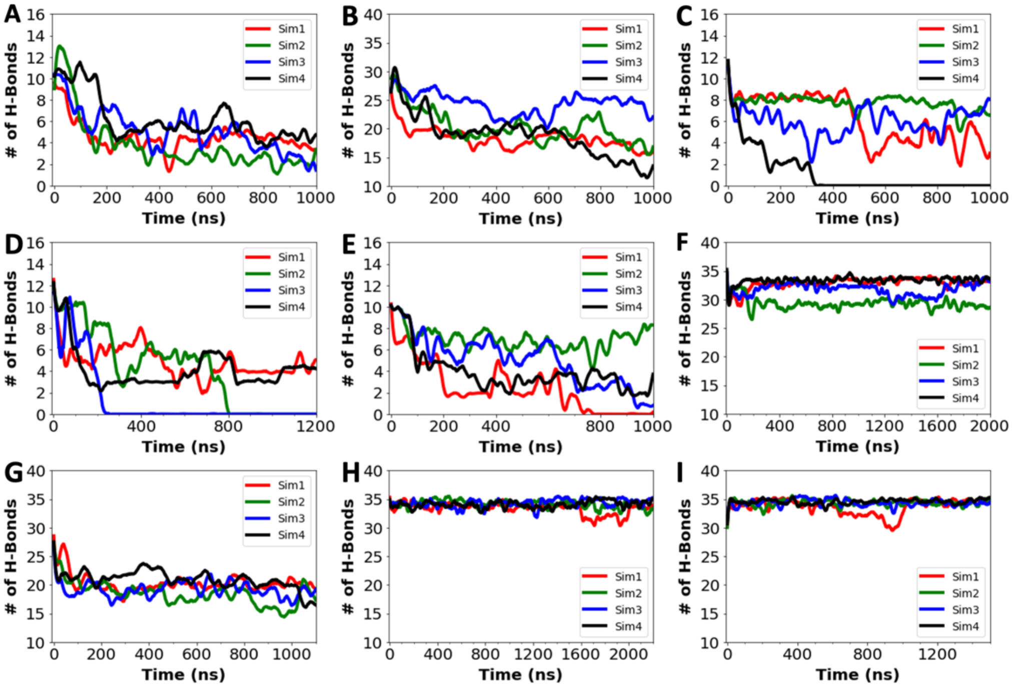


Figure S4. Time traces of the numbers of backbone hydrogen bonds. (A) Aβ parallel N-sheet dimer. (B) Aβ parallel N-sheet tetramer. (C) Aβ42 parallel C-sheet dimer. (D) Aβ42 antiparallel C-sheet dimer. (E) Aβ40 parallel C-sheet dimer. (F) Aβ42 parallel C-sheet tetramer (G) Aβ40 parallel C-sheet tetramer. (H) Aβ42 antiparallel C-sheet tetramer. All the above are in SDS micelles. (H) Aβ42 antiparallel C-sheet tetramer in a DPC micelle.

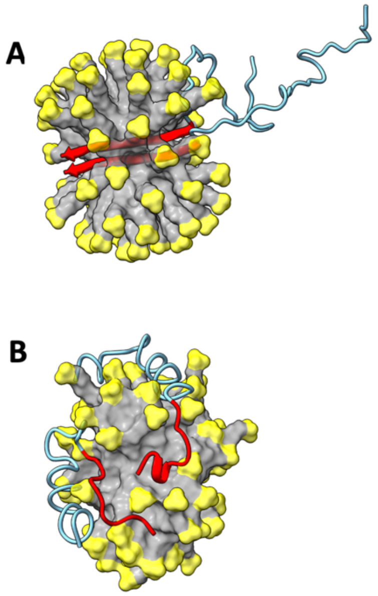

Figure S5. MD simulations of the Aβ40 parallel C-sheet dimer in an SDS micelle. (A) Initial structure. (B) Snapshot at 1000 ns of a simulation. The SDS micelle is shown in surface representation with headgroups in yellow and hydrocarbon tails in grey. Aβ40 molecules are shown in cartoon representation with residues 1-29 in cyan and residues 30-40 in red.

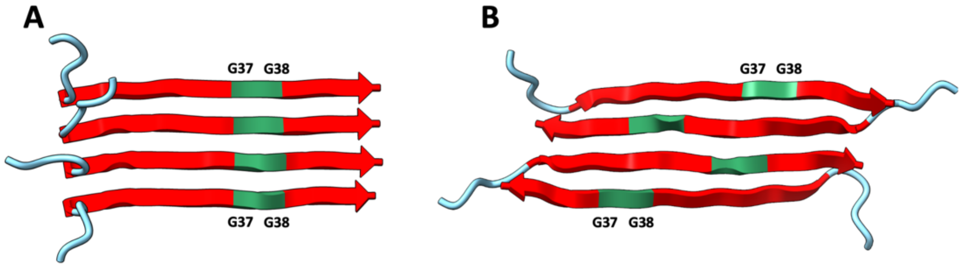


Figure S6. Placements of G37 and G38 in parallel and antiparallel C-sheet tetramers of Aβ42. (A) Parallel C-sheet tetramer. (B) Antiparallel C-sheet tetramer. Aβ42 molecules are shown in cartoon representation with G37 and G38 in green, other residues in the C-strands in red, and the N-terminal residues in cyan.

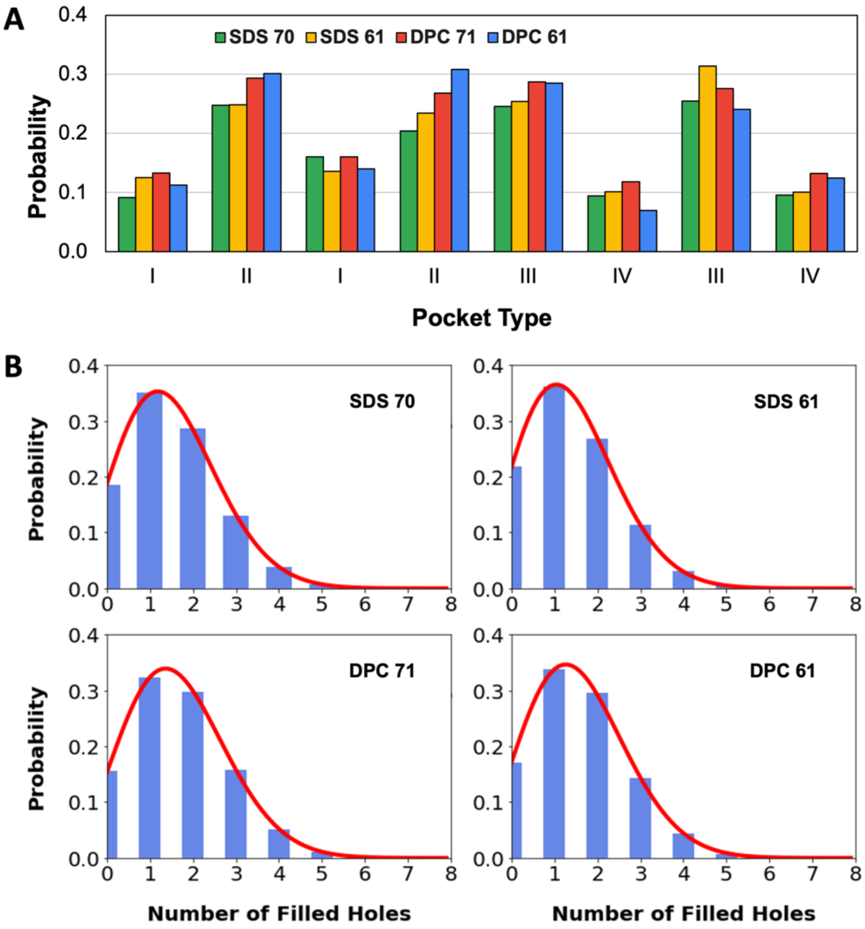


Figure S7. Probabilities of glycine holes being filled by SDS or DPC molecules. (A) Probabilities of individual holes being filled in simulations with SDS and DPC micelles. The legend indicates the detergent molecule and micelle size; e.g., "SDS 70" means a micelle with 70 SDS molecules. (B) Probabilities for a certain number of glycine holes being simultaneously filled at a given time, in four different micelles indicated by the legends. Bars show the raw data; red curves show a binomial distribution, with the probability of a hole being filled given by the average among the eight holes.

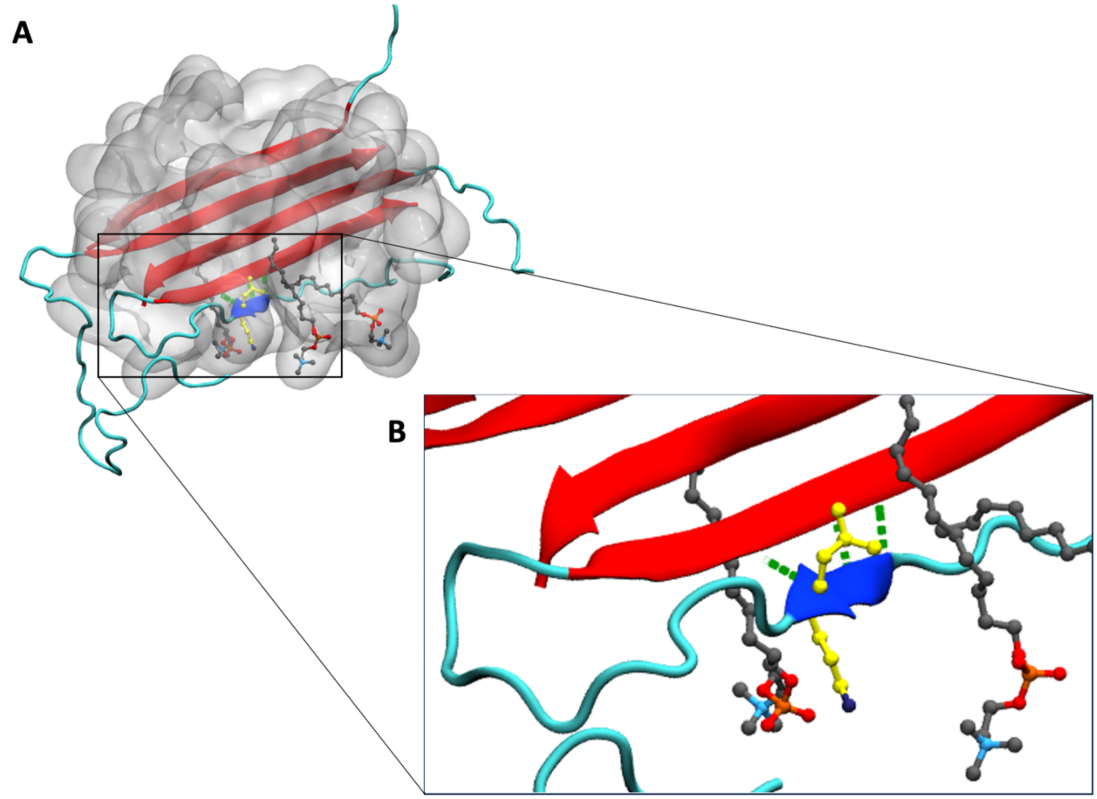


Figure S8. The N-terminal region of an edge chain forms an additional β-strand docked to the antiparallel C-sheet of the Aβ42 tetramer in a micelle with 61 DPC molecules. (A) Snapshot at 162 ns of simulation 3. (B) Zoomed view showing the initiation of the N-strand docking to the edge C-strand. The C-sheet is shown in red; the N-strand is shown in blue; the DPC micelle is shown as a grey surface. Shown in ball-and-stick are the two residues, Lys16 and Leu17, that aid in the N-strand docking (carbon in yellow; nitrogen in dark blue) and the DPC molecules that interact with them (carbon in grey, oxygen in red, phosphorous in orange, nitrogen in light blue). Hydrogen bonds between the edge C-strand and the N-strand are shown as green dashed lines.

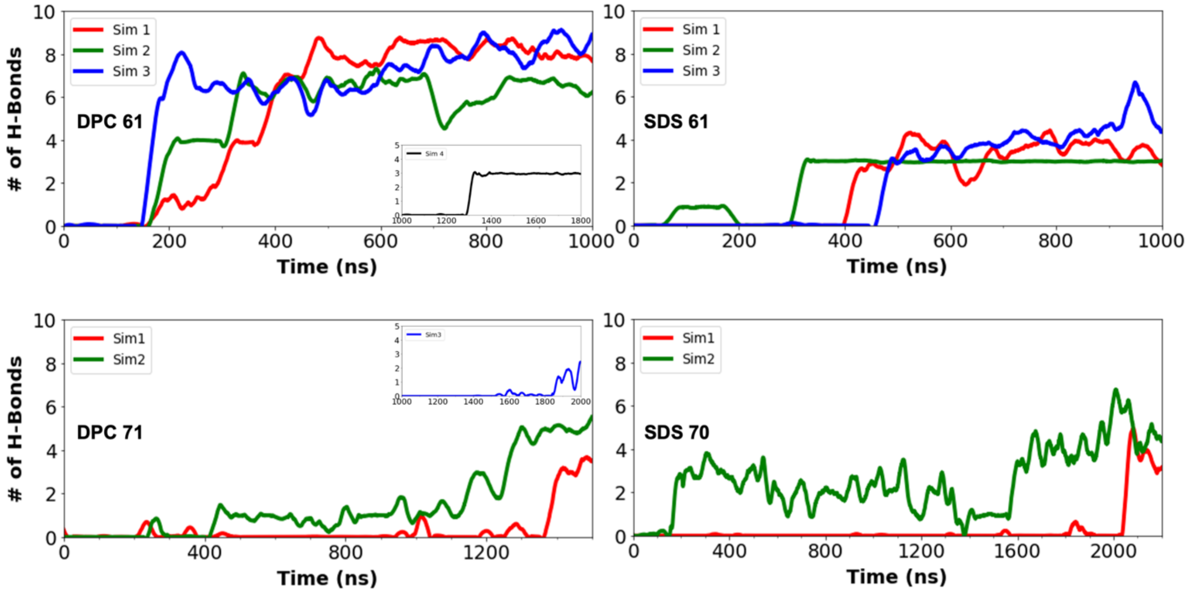


Figure S9. Number of hydrogen bonds between the N-terminal region and the C-sheet, formed in micelles with 61 and 71 DPC molecules or with 61 and 70 SDS molecules.

Movie S1. A 900-ns clip from simulation 1 of the Aβ42 antiparallel C-sheet tetramer in a micelle with 61 DPC molecules. A total of 901 frames were played at 10 frames per second; the 0th frame was the initial structure; the next four frames were from the equilibration stage; and the remaining frames were from the production stage. Frames were separated by 1 ns in simulation time. The C-sheet is shown in red; the N-strand that docks to the C-sheet is shown in blue; an α-helix formed in the N-terminal region of another chain is shown in green; the DPC micelle is shown as a grey surface. Shown in ball-and-stick are the two residues, Lys16 and Leu17, that aid in the N-strand docking (carbon in yellow; nitrogen in dark blue) and the DPC molecules that interact with them (carbon in grey, oxygen in red, phosphorous in orange, nitrogen in light blue). Hydrogen bonds between the edge C-strand and the N-strand are shown as green dashed lines.

Movie S2. A zoomed clip from Movie S1, from 100 ns to 200 ns, highlighting the initiation of N-strand docking to the edge C-strand.
